# The Salivary Microbiome and Predicted Metabolite Production are Associated with Progression from Barrett’s Esophagus to Esophageal Adenocarcinoma

**DOI:** 10.1101/2023.06.27.546733

**Authors:** Quinn S Solfisburg, Federico Baldini, Brittany L Baldwin-Hunter, Harry H Lee, Heekuk Park, Daniel E Freedberg, Charles J Lightdale, Tal Korem, Julian A Abrams

**Affiliations:** Department of Medicine, Boston University School of Medicine, Boston, MA, USA; Program for Mathematical Genomics, Department of Systems Biology, Columbia University Irving Medical Center, New York, NY, USA; Department of Medicine, Columbia University Irving Medical Center, New York, NY, USA; Microbiome and Pathogen Genomics Collaborative Center, Department of Medicine, Columbia University Irving Medical Center, New York, NY, USA; Digestive and Liver Disease Research Center, Columbia University Irving Medical Center, New York, NY, USA; Department of Obstetrics and Gynecology, Columbia University Irving Medical Center, New York, NY, USA; CIFAR Azrieli Global Scholars Program, CIFAR, Toronto, Canada; Herbert Irving Comprehensive Cancer Center, Columbia University Irving Medical Center, New York, NY USA

## Abstract

Esophageal adenocarcinoma (EAC) is rising in incidence and associated with poor survival, and established risk factors do not explain this trend. Microbiome alterations have been associated with progression from the precursor Barrett’s esophagus (BE) to EAC, yet the oral microbiome, tightly linked to the esophageal microbiome and easier to sample, has not been extensively studied in this context. We aimed to assess the relationship between the salivary microbiome and neoplastic progression in BE to identify microbiome-related factors that may drive EAC development. We collected clinical data and oral health and hygiene history and characterized the salivary microbiome from 250 patients with and without BE, including 78 with advanced neoplasia (high grade dysplasia or early adenocarcinoma). We assessed differential relative abundance of taxa by 16S rRNA gene sequencing and associations between microbiome composition and clinical features and used microbiome metabolic modeling to predict metabolite production. We found significant shifts and increased dysbiosis associated with progression to advanced neoplasia, with these associations occurring independent of tooth loss, and the largest shifts were with the genus *Streptococcus*. Microbiome metabolic models predicted significant shifts in the metabolic capacities of the salivary microbiome in patients with advanced neoplasia, including increases in L-lactic acid and decreases in butyric acid and L-tryptophan production. Our results suggest both a mechanistic and predictive role for the oral microbiome in esophageal adenocarcinoma. Further work is warranted to identify the biological significance of these alterations, to validate metabolic shifts, and to determine whether they represent viable therapeutic targets for prevention of progression in BE.

## INTRODUCTION

Esophageal adenocarcinoma (EAC) has seen a dramatic rise in incidence in the past several decades, is often diagnosed at advanced stages, and is associated with poor survival.^1, 2^ The factors that drive EAC remain incompletely understood. Barrett’s esophagus (BE) is the precursor lesion to EAC, but the overwhelming majority of BE patients do not progress to EAC. Established EAC risk factors, including gastroesophageal reflux disease (GERD) and obesity, do not fully explain the rise in its incidence.^3, 4^

Increasing evidence suggests that the microbiome plays an important role in modifying the risk of a variety of epithelial cancers^5–8^ as well as in modulating the response to treatment.^9–11^ Changes in the esophageal microbiome have been observed in EAC and with progression from BE to EAC,^12, 13^ raising the possibility that bacteria contribute to esophageal neoplasia. Reliable sampling of the esophageal microbiome, however, requires invasive procedures. A more accessible “window” to the esophageal ecosystem is the oral microbiome, which was shown to strongly influence it.^14^ A small study of the tumor-associated microbiome in EAC found a high prevalence of domination by oral flora such as *Streptococcus,*^12^ pointing to a link between the oral microbiome and EAC. Oral microbiome alterations have been associated with future risk of EAC^15^, and differences in the oral microbiome of BE patients were described previously in a small study of 49 patients.^16^ Alterations in the oral microbiome have also been associated with poor oral health^17^, which was in itself associated with increased risk of EAC in a recent analysis.^18^ It remains unclear how oral dysbiosis and poor oral health interact in their association with EAC. Finally, little is known with regard to oral microbiome alterations associated with neoplastic progression in BE patients. A clearer understanding of these oral microbiome changes could identify factors that may drive progression of neoplasia, representing novel therapeutic targets.

Here, we profiled the salivary microbiome from 250 patients with various stages of BE and EAC who were undergoing upper endoscopy. We identify multiple characteristics of the oral microbiome associated with neoplastic progression in BE and show that they are independent of oral health. Using metabolic modeling, we predict metabolite profiles associated with alterations in BE, suggesting a mechanistic role for microbially produced metabolites. Finally, we show that the salivary microbiome offers a mild improvement in diagnostic accuracy compared to models based on clinical risk factors. Our results demonstrate the potential of studying the oral microbiome in the context of progression to EAC.

## RESULTS

### Oral microbial composition from a large endoscopy cohort

We recruited 250 adult patients undergoing upper endoscopy and characterized their oral microbiome using 16S rRNA gene sequencing. (**Methods**) A total of 244 patients were included in the analyses: 125 controls without Barrett’s esophagus (BE), and 119 BE patients (20 with non-dysplastic BE, 11 indefinite for dysplasia, 10 low grade dysplasia, 54 high grade dysplasia, and 24 intramucosal (T1a) adenocarcinoma). Patients with BE were more likely to be older (t-test p<0.001), male (Fisher exact p<0.001), white (p=0.001), or ever-smokers (defined as ≥100 lifetime cigarettes smoked) (p=0.003). They were also more likely to have GERD (p<0.001), to be treated with proton pump inhibitor (PPI; p<0.001), to take aspirin (p<0.001), and to have a higher BMI (p<0.001). (**Table 1**) There was no significant difference in the use of mouthwash between BE and non-BE patients (p=0.61), but non-BE patients were more likely to brush their teeth at least daily (98% vs 92%, p=0.03). Compared to non-BE, a significantly higher proportion of patients with BE had tooth loss (63% vs. 42%, p=0.001), largely due to an increase in tooth loss in patients with advanced neoplasia (defined as high grade dysplasia or adenocarcinoma, 66%; non-dysplastic BE, 40%; non-BE, 42%; Fisher’s exact p=0.001). (**Figure 1**) Older age (per year, adjusted OR 1.06, 95% CI 1.04-1.08) and a history of smoking (adjusted OR 2.12, 95% CI 1.07-4.19) were independently associated with tooth loss. (**Supplementary Table 1**) In multivariable analyses adjusting for EAC risk factors (age, male sex, white race, and GERD), tooth loss was associated with a non-significant increased risk of advanced neoplasia (vs. non-BE, adjusted OR 1.49, 95%CI 0.96-2.47). (**Supplementary Table 2**) These results are in line with a recent analyses of data from the Nurses’ Health Study, which found an association between both tooth loss and periodontal disease and risk of esophageal adenocarcinoma, but showed a decrease in association strength after adjusting for covariates.^18^ Established EAC risk factors were independently associated with advanced neoplasia even after controlling for daily tooth brushing, use of mouthwash, and presence of tooth loss, whereas these measures of oral health and hygiene were not independently associated with advanced neoplasia (p=0.21, 0.57, and 0.34, respectively).

**Figure 1.**
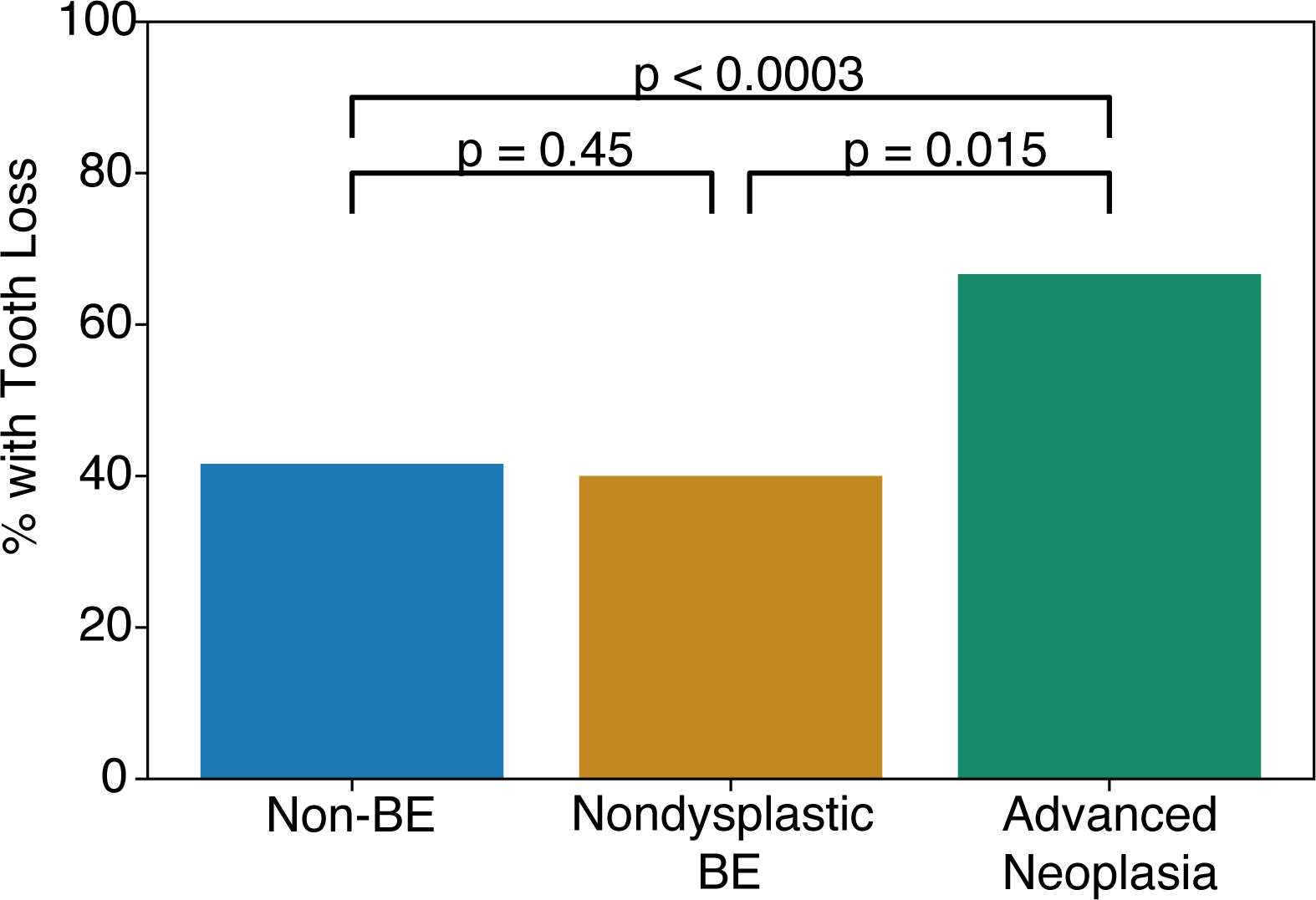
Tooth loss is significantly more common in advanced neoplasia. A significantly higher proportion of patients with advanced neoplasia (high grade dysplasia or esophageal adenocarcinoma) had tooth loss as compared to non-BE and non-dysplastic BE patients combined (Fisher’s exact p = 0.001). P – Fisher’s exact.

**Table 1.** Patient Characteristics. P – t-test or Fisher exact p for difference between BE and non-BE.

|  | All (n=244) | Non-BE<br>(n=125) | BE<br>(n=119) | P-value |
| --- | --- | --- | --- | --- |
| Age, years; mean (SD) | 57.8 (18.7) | 50.9 (18.7) | 65.0 (15.8) | <0.001 |
| Sex, male; N (%) | 140 (57%) | 45 (36%) | 95 (80%) | <0.001 |
| Ever smokers; N (%) | 103 (42%) | 41 (33%) | 62 (52%) | 0.002 |
| BMI mean (SD) | 27.5 (6.7) | 27.0 (7.9) | 28.1 (5.1) | <0.001 |
| Race, white; N (%) | 222 (90%) | 105 (84%) | 117 (98%) | <0.001 |
| GERD; N (%) | 167 (68%) | 64 (51%) | 103 (87%) | <0.001 |
| PPI use; N (%) | 148 (60%) | 41 (33%) | 107 (90%) | <0.001 |
| Aspirin use; N (%) | 77 (31%) | 25 (20%) | 52 (44%) | <0.001 |
| Oral health and hygiene |  |  |  |  |
| Tooth loss; N (%) | 127 (52%) | 52 (42%) | 75 (63%) | <0.001 |
| Tooth brushing frequency, ≥ daily; N (%) | 233 (95%) | 123 (98%) | 110 (92%) | 0.03 |
| Mouthwash frequency, ≥ daily; N (%) | 139 (56%) | 69 (55%) | 70 (59%) | 0.61 |

### The oral microbiome of BE patients is progressively altered with dysplastic changes

To assess whether the salivary microbiome is associated with neoplastic progression, we focused our analyses on comparisons between three groups: non-BE (n=125), non-dysplastic BE (n=20), and advanced neoplasia (high grade dysplasia and adenocarcinoma [HGD/EAC]; n=78). We found that neoplastic progression was associated with significantly lower alpha diversity (Shannon: Kruskal-Wallis p=0.005, **Figure 2A**; Simpson: p=0.0029, **Supplementary Figure 1**). Compared to patients without BE, the alterations in alpha diversity were more pronounced in patients with advanced neoplasia than in those with nondysplastic BE (non-BE vs. non-dysplastic BE, Mann-Whitney p=0.11; non-BE vs advanced neoplasia, p=0.0006). There was no significant difference in alpha diversity comparing nondysplastic BE with advanced neoplasia (p=0.23). We further found that the oral microbiome from patients with advanced neoplasia tended to cluster separately than the rest of the cohort (weighted UniFrac, ANOSIM p<0.001; **Figure 2B**). Similar results were found when including all the subjects in their individual groups (non-BE, nondysplastic BE, IND, LGD, HGD, and EAC; **Supplementary Figure 2**). Our results indicate that the salivary microbiome alterations observed in advanced neoplasia are reflected both in the diversity of each individual’s microbiome (alpha diversity) and in compositional differences between individuals (beta diversity).

**Figure 2.**
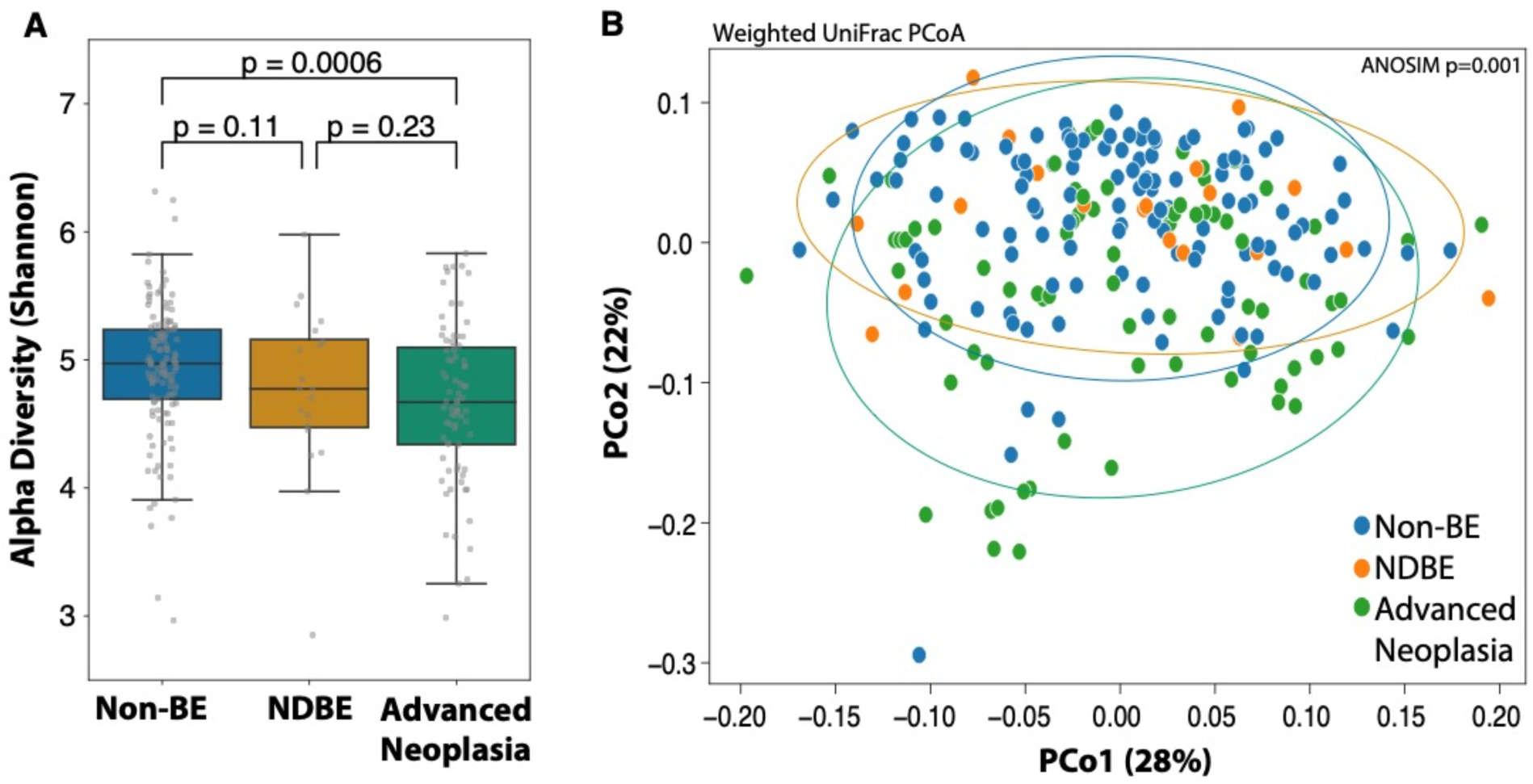
A microbial signature of BE. (A) Patients with advanced neoplasia have significantly reduced alpha diversity compared to non-BE patients. Kruskal-Wallis overall p-value=0.005. p – Mann-Whitney U test. (B) Weighted UniFrac PCoA demonstrated significant clustering of patients with advanced neoplasia (ANOSIM p=0.001). NDBE, non-dysplastic BE; advanced neoplasia – high grade dysplasia or esophageal adenocarcinoma; ellipse, 2 standard deviation sigma ellipse.

We next checked whether specific microbes were associated with BE progression. We therefore compared relative abundance of different OTUs between all BE patients vs. non-BE using ALDEx2.^19^ (**Methods**) A total of 26 OTUs were identified as differentially abundant (p < 0.05, FDR corrected at 0.1; **Figure 3**). To assess whether the dysbiotic signature associated with BE is more pronounced with dysplastic changes, we checked whether these 26 OTUs were correlated with progression across the neoplastic spectrum, from no dysplasia to EAC (Methods). There was a significant association (p<0.05, FDR corrected at 0.1) for 23 of the 26 taxa, with a clear shift in composition with neoplastic progression, notably in the transition from low grade dysplasia (LGD) to high grade dysplasia (HGD). (**Figure 3**) This transition in composition from LGD to HGD is consistent with our prior observations of esophageal microbiome alterations with progression to EAC.^20^ The taxonomic alterations associated with progression were notable for increased relative abundance of several *Streptococcus* species. Streptococci form biofilms in the oral cavity^21–23^.

**Figure 3.**
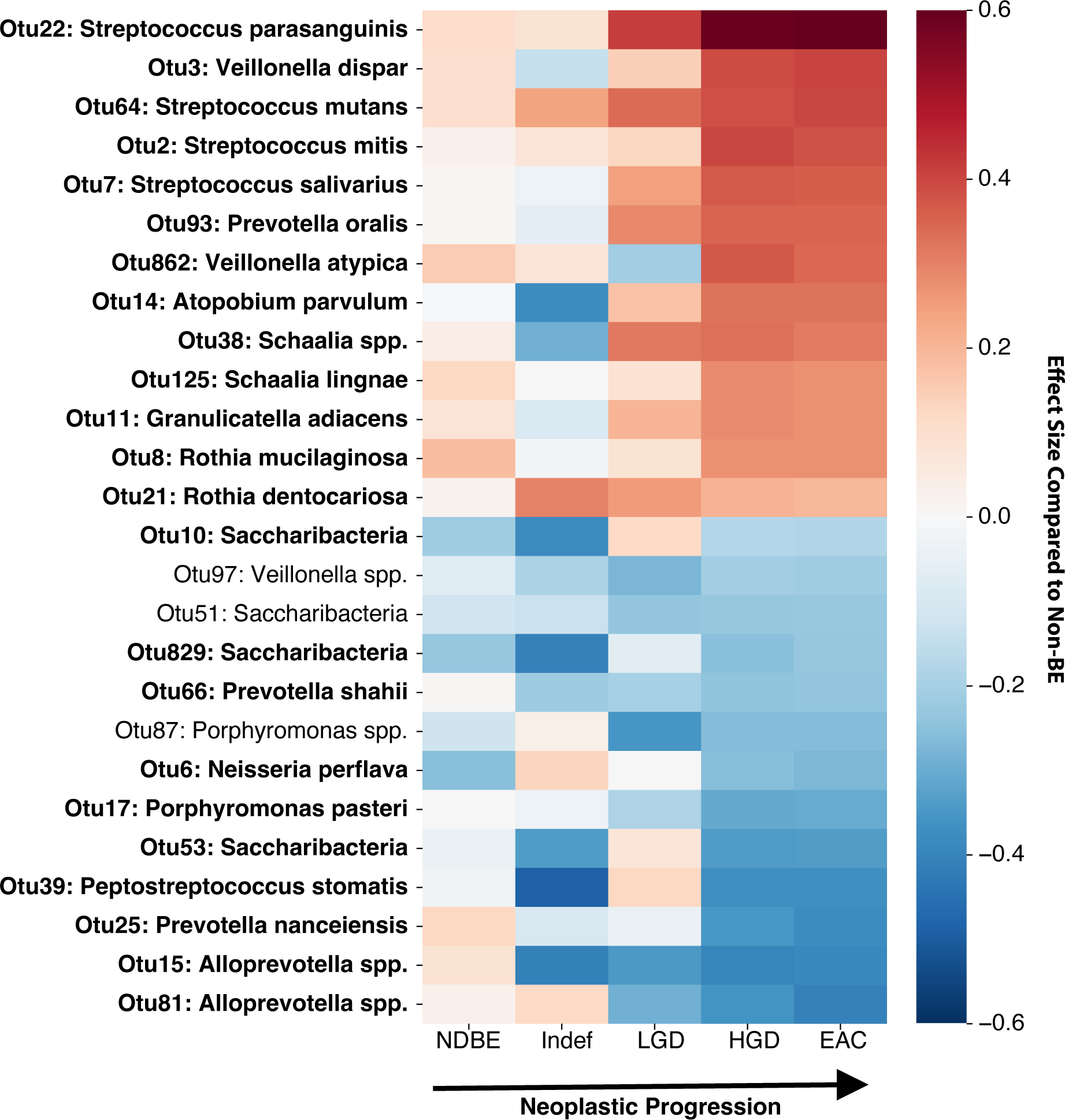
Increased oral dysbiosis with progressive dysplasia. Shifts in the oral microbiome compared to non-BE patients (shown as ALDEx2 effect sizes) were more pronounced with progression from no dysplasia to EAC, particularly notable in patients with high grade dysplasia and EAC. Bolded OTUs were significantly associated with neoplastic progression. (p < 0.05, FDR corrected at 0.1; **Methods**) NDBE, nondysplastic Barrett’s esophagus; Indef, indefinite for dysplasia; LGD, low grade dysplasia; HGD, high grade dysplasia; EAC, esophageal adenocarcinoma.

### The salivary microbiome is associated with advanced neoplasia even when controlling for tooth loss

Tooth loss is known to be strongly associated with oral microbiome composition^17^, and there was an increased proportion of advanced neoplasia patients with tooth loss. (**Figure 1**) Comparing patients who did and did not have all or most of their natural adult teeth, we found that those with tooth loss had lower alpha diversity (Shannon, Mann-Whitney p=0.001; **Supplementary Figure 3A**), and that oral microbiomes from both groups clustered separately (weighted UniFrac, ANOSIM p=0.003; **Supplementary Figure 3B**). We also identified 29 OTUs that had significantly different abundance between patients with and without tooth loss. (ALDEx2 p < 0.05, FDR corrected at 0.1; **Figure 4**) As poor oral health is associated with oral dysbiosis, this raised the question of whether the oral microbiome is associated with advanced neoplasia independent of tooth loss.

**Figure 4.**
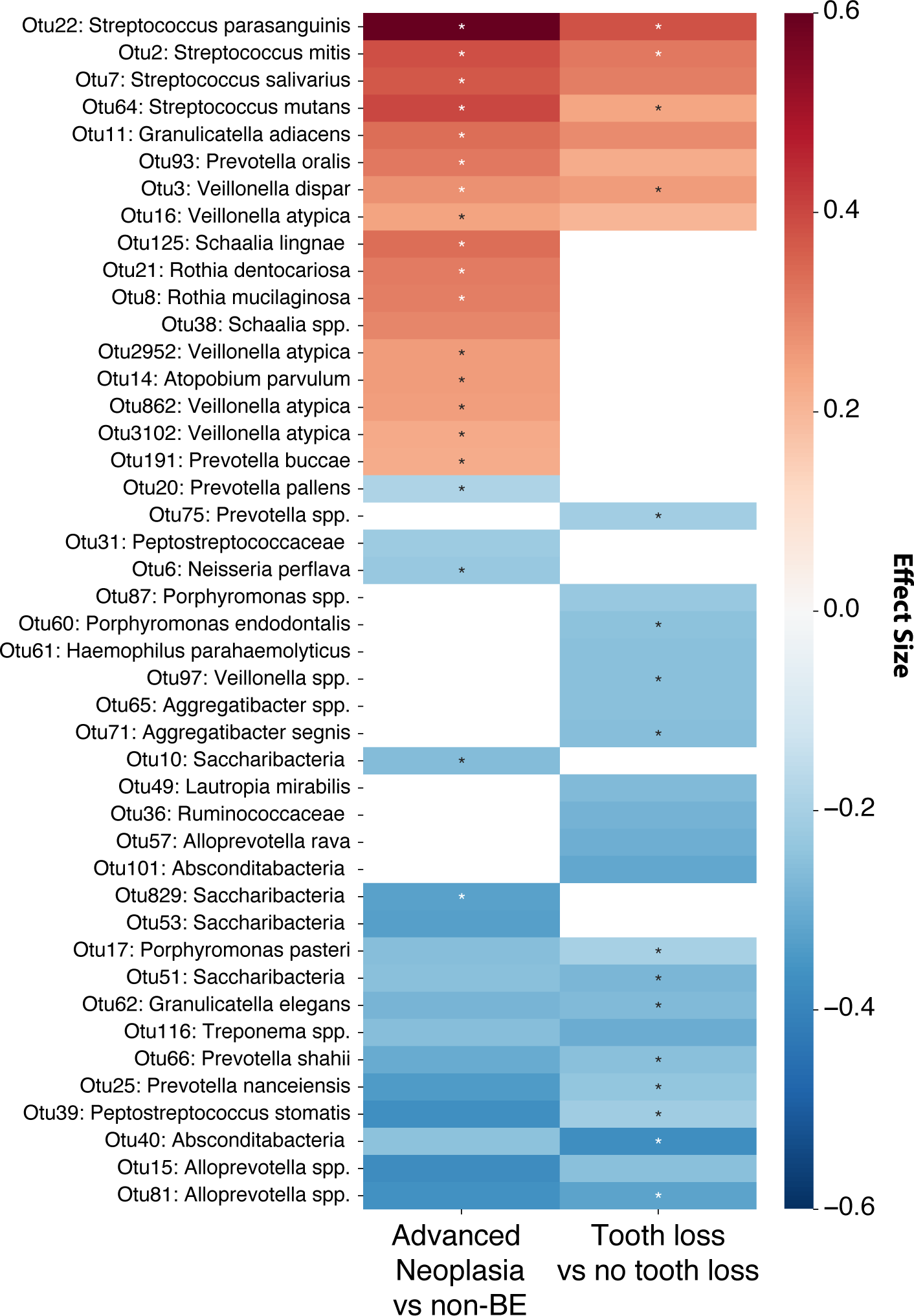
Oral microbes are independently associated with advanced neoplasia and tooth loss. Differentially abundant taxa in patients with advanced neoplasia (high-grade dysplasia or esophageal adenocarcinoma) compared to non-BE (left) and in patients with and without tooth loss (right). Many taxa were associated with both neoplasia and tooth loss, yet most of these remained significantly differentially abundant after adjusting for tooth loss and advanced neoplasia, respectively (ALDEx2 p < 0.05, FDR corrected at 0.1; denoted by asterisks).

We first examined whether salivary microbiome composition as a whole is associated with advanced neoplasia independently of tooth loss. We therefore calculated microbiome principal coordinates (PCos) using weighted UniFrac distances and used the top five PCos, which represented two thirds of the variance in microbiome composition. We then used multivariable logistic regression and found that PCo2 (explaining 22% of microbiome variation) and PCo4 (4.6%) were independently associated with advanced neoplasia. (p<0.001 and p=0.004, respectively) We then added the major EAC risk factors (age, sex, race, BMI, GERD history, and smoking history) to the model, and found that PCo2 remained independently associated with advanced neoplasia (p=0.004), suggesting that salivary microbiome composition represents a potential novel independent risk factor for EAC. Adding tooth loss to the model did not alter the association between PCo2 and advanced neoplasia (p=0.004), and in this model tooth loss was not independently associated with advanced neoplasia (p=0.12). Our results suggest that the association of tooth loss with advanced neoplasia is mediated through the oral microbiome.

We next assessed whether associations between specific oral taxa and advanced neoplasia are independent of tooth loss. Of the 33 taxa associated with advanced neoplasia (N=78) vs. non-BE (N=125; ALDEx2 p < 0.05; FDR corrected at 0.1), 18 were also associated with tooth loss. (**Figure 4**) After adjusting for tooth loss in a generalized linear model, 20 of these taxa remained significantly associated with advanced neoplasia. (ALDEx2 p < 0.05, FDR corrected at 0.1) Notably, the four OTUs with the greatest increase in relative abundance in advanced neoplasia were all assigned to the genus *Streptococcus*, and the increased abundance of these *Streptococcus* OTUs in advanced neoplasia was independent of tooth loss. This corresponds with previous studies that found that the tumor-associated microbiome in EAC is often dominated by *Streptococcus* species.^12^

### Metabolic modeling predicts distinct metabolic secretion capabilities in advanced neoplasia

Metabolite production by microbial communities is an important modality by which the microbiome affects the host. In order to assess if microbially produced metabolites might play a role as a driver or biomarker of neoplasia, we used microbiome community-scale metabolic models to predict metabolite secretion by the microbiome for every sample. (**Methods**) We found significant clustering of predicted metabolite profiles comparing advanced neoplasia cases with non-BE subjects (PERMANOVA p=0.001). Using principal components analysis, we found notable shift in the second component (15% explained variance; Mann-Whitney p=0.0003). (**Figure 5A**) Forty-four predicted metabolites had significantly altered abundance (p < 0.05, FDR corrected at 0.1) in advanced neoplasia. (**Figure 5B**) Notable alterations included increased predicted levels of L-lactic acid (p=0.023), a by-product of aerobic glycolysis, a hallmark of cancer which can contribute to neoplasticity;^24^ and 2-ketobutyric acid (p=0.033), previously reported to support mitochondrial respiration and cell proliferation.^25^ We also predicted that advanced neoplasia features a decrease in butyric acid (p=0.0089), a key promoter of gut homeostasis that was previously shown to be depleted in colon cancer and inflammatory bowel disease;^26–28^ and a decrease in L-tryptophan (p=0.0017). (**Figure 5C**) Circulating levels of tryptophan have been inversely associated with colon cancer risk^29^, and melatonin, a by-product of L-tryptophan metabolism, is under investigation for EAC prevention.^30^ The pattern of the shifts we predict is therefore consistent with the potential promotion of proliferation, inflammation, and cancer.

**Figure 5.**
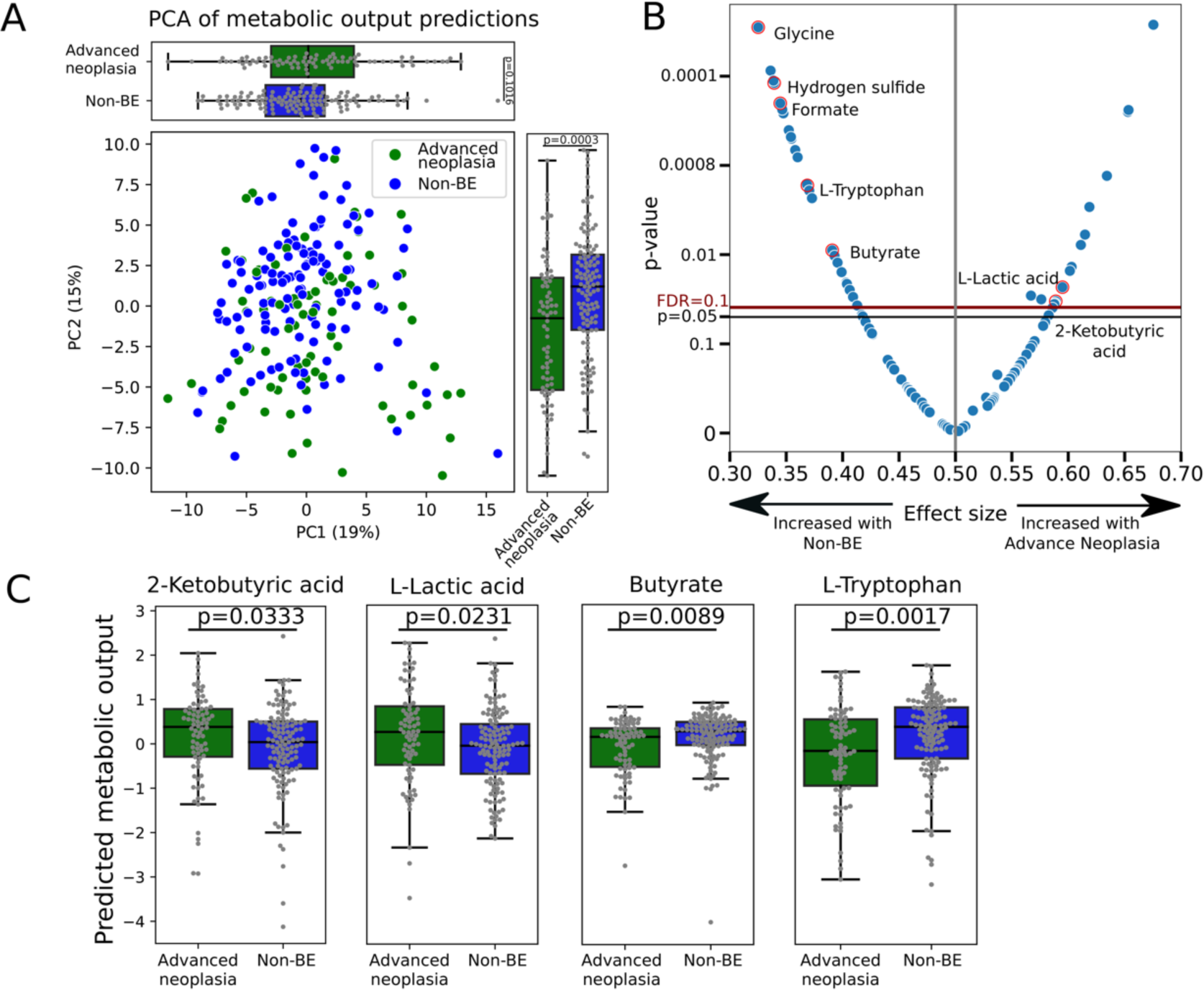
The predicted metabolic profile is altered in patients with advanced neoplasia. (A) Significant clustering by advanced neoplasia status on principal components analysis (PERMANOVA p=0.001), with pronounced shifts in PC2 (p=0.0003). (B) Volcano plot demonstrating differentially abundant metabolites in advanced neoplasia. (C) Significantly altered predicted levels of L-lactic acid, 2-ketobutyric acid, butyric acid, and L-tryptophan in advanced neoplasia. Plot capped at −4 for butyrate. P, Mann-Whitney test

### Salivary microbiome data improves on clinical risk factors-based prediction of advanced neoplasia

We next performed an exploratory analysis to determine whether salivary microbiome features in this cohort could be used to distinguish advanced neoplasia from non-BE patients. As a baseline, we first trained a gradient boosted decision trees model which uses clinical risk factors for EAC and tooth loss to classify advanced neoplasia. The classifier was tested in cross-validation on patients not seen in the training of that model and achieved an area under the receiver operating characteristic curve (AUROC) of 0.84 (95%CI 0.79-0.89). The same process was then used to train a classifier using microbiome data, whereas, within each training fold, 10 OTUs were selected based on a Kruskal-Wallis test. This classifier had an AUROC of 0.72 (95%CI 0.65-0.79). Finally, a model trained on the combination of both microbiome data and EAC risk factors resulted in somewhat higher model accuracy, producing an AUROC of 0.88 (95%CI 0.83-0.91; vs. clinical risk factor model AUROC 0.84, 95%CI 0.79-0.89; DeLong p=0.053). (**Supplementary Figure 4**) A combined microbiome and clinical risk factor model with the outcome limited to HGD and excluding intramucosal EAC showed similar results with an AUROC 0.86 (95%CI 0.80-0.92), compared to an AUROC of 0.83 (95%CI 0.77-0.89) using only clinical risk factors for the same task. (**Supplementary Figure 5**)

## DISCUSSION

In this cross-sectional study of patients with and without BE, we detected marked shifts in the salivary microbiome with progression to EAC, with changes that appeared to be most pronounced in patients with advanced neoplasia. These changes included reduced diversity as well as significantly increased relative abundance of several taxa in the genus *Streptococcus*. As in previous studies, we found that tooth loss is more common in patients with advanced neoplasia. However, we show that many of the salivary microbiome associations observed in BE and advanced neoplasia persisted even when accounting for it. Further, we used metabolic modeling to identify distinct predicted metabolic secretion capabilities in advanced neoplasia.

Our findings add to the growing body of evidence that the oral microbiome is linked to the esophageal microbiome and may contribute to esophageal neoplasia. In a case-control study of patients with EAC, BE, and controls, the EAC-associated microbiome had significantly reduced alpha diversity, similar to our observations in saliva.^12^ Interestingly, in that study 5/15 of the EAC tumors were dominated by *Streptococcus* spp. (relative abundance 69%-98%). In our salivary microbiome analyses, 4 of the 5 taxa most strongly associated with advanced neoplasia were also *Streptococcus* spp. Our group conducted a randomized controlled trial and found that an antimicrobial mouth rinse can produce esophageal microbiome and tissue gene expression changes, highlighting the relevance of oral bacteria to esophageal disease.^31^ In another cohort study analyzing mouth rinse samples from patients enrolled in two large cancer prevention studies, oral microbiome alterations were noted to precede an EAC diagnosis by several years.^15^ In a small study of 49 patients we previously noted marked salivary microbiome alterations associated with BE and also with advanced neoplasia.^16^

Our study features the use of microbiome metabolic models to identify broad shifts in metabolites predicted to be produced by the saliva microbiome. Many of the predicted changes to metabolite outputs correspond with existing knowledge. Lactic acid, for example, was predicted to be increased in advanced neoplasia. Lactic acid can serve as a major energy source for proliferative cancer cells, and is known to activate hypoxia inducible factors, which in turn contribute to proliferation, angiogenesis, and other neoplastic features.^24^ Our findings could therefore support the hypothesis that the oral and esophageal microbiota promotes EAC development and progression via production of metabolites.^32^ Our findings further correspond with a previous study detecting lactic-acid bacteria in many esophageal adenocarcinomas.^12^ However, the biological significance of predicted metabolite production is unclear, and future studies are needed to validate these predictions and to elucidate the biological effects of specific bacterial metabolites on esophageal neoplasia.

Prior work has associated tooth loss and periodontal disease with increased risks of esophageal squamous cell cancer and gastric cancer, and a recent study found an association between both tooth loss and periodontal disease and risk of esophageal adenocarcinoma.^18^ This indicates a potential confounding effect, as tooth loss is also associated with major alterations in oral microbiome composition.^17^ Our study offers an explanation for these associations, demonstrating that salivary microbiome composition is independently associated with advanced neoplasia, even when adjusting for EAC risk factors and for tooth loss. These findings suggest that the association between tooth loss and esophageal neoplasia is mediated by changes in the salivary microbiome, and that the salivary microbiome may represent a novel independent risk factor for EAC.

We performed exploratory analyses to assess whether the salivary microbiome could discriminate patients at highest EAC risk. The salivary microbiome is highly suitable for diagnostics, as it is stable over time^33–35^, especially compared to other body sites^36^, and is resistant to perturbations.^37^ Addition of a microbiome-based classifier to EAC risk factors resulted in modest improvement in discrimination. However, the current study was not specifically designed to address this question, and future studies should explore further the salivary microbiome as a potential biomarker for advanced neoplasia.

Important strengths of the current study include the relatively large sample size and the inclusion of oral health and hygiene information from patients. The large sample size allowed for the detection of significant microbiome alterations, even when correcting for multiple comparisons. Previous studies of the oral microbiome in BE and EAC have not included oral health and hygiene data, key potential confounders. The patients were well characterized, with data collected on key EAC risk factors including GERD history, BMI, and smoking, which permitted microbiome analyses adjusting for these variables. The BE patients in the study were demographically similar to BE populations from other studies, enhancing the generalizability of the findings. Lastly, novel methods for predicted microbiome metabolic profiling allowed for insights into functional correlates of the salivary microbiome alterations.

The study does have certain limitations. There were a relatively small number of non-dysplastic BE patients, limiting analyses in this subgroup. Analyses did not incorporate dietary intake; however, previous studies suggest that diet has minimal impact on salivary microbiome composition.^38–40^ No conclusions can be drawn with regard to temporality in this cross-sectional study. It is possible that the observed salivary microbiome alterations were caused by BE-associated advanced neoplasia, although we believe that this is unlikely. Tooth loss was self-reported rather than measured, and periodontal disease was not directly assessed. Community-scale metabolic models also have notable limitations. Our analysis was based on 16S rRNA gene sequencing, which does not allow us to tailor models to specific strains or genetic potential present in each sample. Additionally, while genome-scale models have been curated for common gut commensals, to our knowledge, such efforts have not been done for oral microbes. Consequently, some models may be missing, while existing ones may lack representation of niche-specific metabolic capacity. Despite these limitations, these models allow a systematic application of biochemical and genetic knowledge to our analysis and raise interesting hypotheses that could be experimentally validated.

In conclusion, patients with BE-associated advanced neoplasia have a markedly altered salivary microbiome, and analyses of taxonomic alterations associated with stages of progression from BE to EAC appear to indicate that these changes are most notable at the transition from low- to high-grade dysplasia. Increased tooth loss was also observed with progression to EAC, although the salivary microbiome alterations were largely independent of tooth loss, suggesting that the association of tooth loss with advanced neoplasia is mediated through the oral microbiome. There were marked increases in various taxa in the genus *Streptococcus* in advanced neoplasia, possibly pointing to a biological contribution of these bacteria to neoplastic progression. In addition to the microbiome alterations, progression to EAC was associated with numerous changes to predicted bacterial metabolite production, with notable alterations that suggest possible proneoplastic effects related to these shifts. Further work is warranted to identify the biological significance of the microbiome alterations, to validate metabolic shifts, and to determine whether they represent viable therapeutic targets for prevention of progression in BE.

## METHODS

### Study Design

A total of 250 patients with and without BE undergoing upper endoscopy at Columbia University Irving Medical Center (New York, NY) were prospectively enrolled from February 2018 through February 2019. Patients were ≥18 years old and scheduled to undergo endoscopy for clinical indications. Patients were excluded if they had a concurrently scheduled colonoscopy, had a history of gastric or esophageal surgery, a history of esophageal squamous cell cancer, or use of antibiotics, steroids, or other immunosuppressants in the 3 months prior to the procedure. This study was approved by the Columbia University Institutional Review Board. All patients provided written informed consent.

Data were collected on patient demographics and anthropometrics (to calculate BMI) as well as clinical information including medical history, history of gastro-esophageal reflux disease (GERD; defined as experiencing frequent heartburn or fluid regurgitation), medication use at time of enrollment (with specific notation of daily use of proton pump inhibitors (PPIs), histamine-2 receptor antagonists, statins, and daily use of aspirin and non-steroidal anti-inflammatory drugs), alcohol history, and smoking history (ever smoking defined as having smoked >100 lifetime cigarettes). Data were collected on self-reported oral health and hygiene. Tooth loss was assessed using categories adapted from Borningen et al.^17^: all or most of natural adult teeth, partial plates or implants, full upper dentures or implants, full lower dentures or implants, full upper and lower dentures or implants. Data were also collected on tooth brushing and mouthwash use.

Patients did not eat or drink after midnight prior to the endoscopy and saliva collection; saliva was collected prior to the endoscopy. Patients were categorized as BE if they had a history of endoscopically suspected BE with intestinal metaplasia on esophageal biopsies. BE patients were further categorized based on the highest degree of neoplasia ever (no dysplasia (NDBE), indefinite for dysplasia (IND), low grade dysplasia (LGD), high grade dysplasia (HGD), adenocarcinoma (EAC)).

### Microbiome Sequencing and Analysis

The 16S rRNA V3-V4 region was amplified using Illumina adapter-ligated primers.^41^ The Illumina Nextera XT v2 index sets A-D were used to barcode sequencing libraries. Libraries were sequenced on an Illumina MiSeq using the v3 reagent kit (600 cycles) and a loading concentration of 12 pM with 10% PhiX spike-in. Sequences were assigned to operational taxonomic units (OTUs) using USEARCH^42^ with ≥ 97% sequence homology. Taxonomic assignments for the OTUs were based on the Human Oral Microbiome Database (HOMD).^43^ Any subsequently unassigned OTUs were assigned by referencing the Ribosomal Database Project (RDP).^44^ Samples were subsampled to 10,000 reads to compare across even sequencing depths while minimizing data loss. Five patients were excluded after sequencing because of relatively low sequencing depth with <10,000 total reads per sample. The median read count for the full cohort was >33,000. One patient was excluded because of a history of both EAC and esophageal squamous cell carcinoma.

### Microbiome metabolic modeling of oral microbial communities

Microbiome metabolic modeling was performed using the Microbiome Modeling Toolbox (COBRA toolbox commit: 71c117305231f77a0292856e292b95ab32040711) ^45, 46^ and the AGORA metabolic models (AGORA 1.02).^47^ All computations were performed in MATLAB version 2019a (Mathworks, Inc.), using the IBM CPLEX (IBM, Inc.) solver. We first matched species detected by our microbial sequencing analysis with the ones present in AGORA.^47^ Because AGORA metabolic models are available at the strain level, we generated species-level models using the createPanModels.m function of the Microbiome Modeling Toolbox (MMT)^45^ as previously described.^48^ To increase the number of species represented in our microbiome models we chose genus-level representative models for abundant microbes present in the oral cavity with >5% relative abundance in more than 10 samples. There were six species without a corresponding metabolic model, and these were either grouped with similar species or excluded from the analyses (See **Supplementary Table 3** for details).

We then used the mgPipe.m automated pipeline of the MMT to build and interrogate sample-specific microbiome metabolic models. Briefly, for each sample, personalized microbiome models are created by joining species-level metabolic models using the compartmentalization technique^49^; a lumen compartment enabling microbial metabolic interactions is added, as well as additional input and output compartments, allowing microbiome intake and secretion of metabolites. Altogether our microbiome models included 160 microbial species with an average of 50 species for each sample and a maximum of 69. As constraint-based metabolic modeling benefits from a specification of the metabolic environment such as media and carbon source availability^50^, we applied a “western diet”^51^ to each sample in the form of constraints on the metabolites uptake reactions.^51^ Finally, to obtain metabolic predictions, we used the Net Maximal Production Capabilities (NMPCs) through the mgPipe pipeline^45^ to provide predictions of the metabolite secretion profile of each sample. To detect significant changes in NMPCs distributions between cases and controls a Mann-Whitney U test was performed for each retained NMPCs. Only NMPCs which were present in at least 10% of the cases and had at least a value of 0.01 were retained for the significance analysis. FDR correction using the Benjamini–Hochberg procedure was applied.

### Statistical Analysis

The primary groups of comparison were BE patients with advanced neoplasia (HGD or EAC), non-dysplastic BE (NDBE) and non-BE controls. Grouping high grade dysplasia and intramucosal adenocarcinoma together as advanced neoplasia reflects common practice as well as clinical guidelines for treatment.^52^ There is extremely low inter-observer agreement (even among expert gastrointestinal pathologists) for the diagnosis of LGD^53–55^, as inflammation-induced cytologic atypia mimics the findings of LGD. As a result, while estimates of cancer risk for LGD are relatively low on average,^54^ these estimates vary widely, thus making interpretations of findings for this group challenging. Patients with low grade dysplasia or indefinite for dysplasia were included in analyses assessing for alterations in the oral microbiome across the entire BE neoplastic spectrum. While there was no *a priori* reason to suspect that endoscopic therapy would have altered the salivary microbiome, comparisons were made between those patients with LGD or worse who had (n=78) and had not (n=10) received prior endoscopic therapy. There were no differences in alpha diversity (p=0.16), no evidence of clustering on beta diversity analyses (ANOSIM p=0.13), and no differentially abundant taxa. Thus, treated and untreated patients were grouped together for all analyses.

Categorical variables were compared across groups using Fisher’s exact tests. Continuous variables were analyzed using t-tests or rank sum tests as appropriate, with ANOVA and Kruskal Wallis tests for ≥3 groups. For purposes of analyses, tooth loss was dichotomized as having all or most of natural adult teeth (yes/no). Multivariable logistic regression was performed to assess the association between tooth loss and advanced neoplasia, adjusted for known EAC risk factors (age, sex, GERD, body mass index (BMI), smoking).

Alpha diversity was evaluated using the Shannon diversity index and beta diversity using weighted UniFrac^56^ distances. Groups were compared using both permutational multivariate analysis of variance (PERMANOVA) for predicted metabolite profiles and analysis of similarities (ANOSIM) for microbial compositions. To find differential abundances between study groups, the ALDEx2^19^ R package was used. For differential abundance analyses, only OTUs present in at least 5% of all samples were included to allow for more meaningful comparisons. ALDEx2 was used to compare worst histological grades of BE as an ordinal variable in a generalized linear model and to assess correlation of BE-associated OTUs with neoplastic progression using aldex.corr to treat worst histological grade as a continuous variable. ALDEx2 was also used to find significance for differentially abundant taxa in a multivariate model with both advanced neoplasia and tooth loss.

Generalized linear models were used to assess differential relative abundance of bacterial taxa in advanced neoplasia, adjusted for tooth loss. Multivariable logistic regression was performed to detect associations between advanced neoplasia and microbiome composition (represented by its top five principal coordinates), adjusted for EAC risk factors (age, sex, race, BMI, smoking, GERD). Supervised machine learning was used to classify patients with advanced neoplasia using the LightGBM package.^57^ Three models were created: 1) EAC risk factors alone (age, sex, race, BMI, smoking, GERD); 2) microbiome features alone; and 3) EAC risk factors and microbiome features together. Model parameters were optimized per fold in 10-fold cross-validation, with strict train-test sterility. The output of the models were predicted probabilities of whether a patient has advanced neoplasia or no BE, with the goal of identifying the patients at highest risk of mortality from EAC.

All statistical analyses were performed in Python or R. Statistical significance was defined as p<0.05. Differential abundance analyses were corrected for multiple comparisons using the Benjamini-Hochberg procedure, and corrected statistical significance was defined as p<0.1. 95% confidence intervals for AUCs were calculated using the DeLong method using pROC.^58^

## Data Availability

16S rRNA gene sequencing files were uploaded to NCBI Sequence Read Archive (PRJNA785879).

## DECLARATIONS

### Ethical approval and consent to participate

This study was approved by the Columbia University Institutional Review Board. All patients provided written informed consent.

### Consent for publication

Not applicable

### Availability of data and materials

16S rRNA gene sequencing files were uploaded to NCBI Sequence Read Archive (PRJNA785879).

### Authors’ contributions

Quinn Solfisburg performed data analysis, data interpretation, and original drafting of the manuscript. Federico Baldini performed data analysis, data interpretation, and assisted with drafting the manuscript. Brittany Baldwin Hunter performed study conduct and provided critical input to the manuscript. Harry Lee performed data analysis and provided critical input to the manuscript. Daniel Freedberg performed data interpretation and provided critical input to the manuscript. Charles Lightdale performed study conduct and provided critical input to the manuscript. Tal Korem supervised data analysis, performed data interpretation, and assisted with drafting the manuscript. Julian Abrams designed the study, supervised data analysis, performed data interpretation, and assisted with drafting the manuscript. All authors approved the final version of the manuscript.

## Acknowledgments

None

