## Supplementary Information for "The Salivary Microbiome and Predicted Metabolite Production are Associated with Progression from Barrett’s Esophagus to Esophageal Adenocarcinoma"

Supplementary Materials

**Supplementary Table 1.** Multivariable Logistic Regression Analyses Assessing Associations Between EAC Risk Factors and Tooth Loss.

| | Odds Ratio | 95% CI |
|---|---|---|
| Male Sex | 0.85 | 0.43-1.67 |
| White Race | 1.48 | 0.51-4.31 |
| GERD | 0.86 | 0.42-1.79 |
| Age (year) | 1.06 | 1.04-1.08 |
| Ever Smoking | 2.12 | 1.07-4.19 |
| BMI >25 | 1.41 | 0.94-2.10 |

**Supplementary Table 2.** Multivariable Logistic Regression Analyses Assessing Association Between Tooth Loss and Advanced Neoplasia.

| | Odds Ratio | 95% CI |
|---|---|---|
| Tooth loss | 1.49 | 0.96-2.47 |
| Male Sex | 5.80 | 2.63-12.8 |
| White Race | 13.2 | 1.55-111 |
| GERD | 6.51 | 2.65-16.0 |
| Age (year) | 1.04 | 1.01-1.06 |
| Ever Smoking | 1.57 | 0.73-3.40 |
| BMI >25 | 1.54 | 0.66-3.38 |

**Supplementary Table 3.** Species without a corresponding metabolic model in the predicted metabolite analyses.

| Species: | Grouped with: |
|---|---|
| *Porphyromonas pasteri* | *Porphyromonas gingivalis* |
| *Neisseria sicca* | *Neisseria subflava* |
| *Neisseria perflava* | *Neisseria subflava* |
| *Prevotella histicola* | *Prevotella melaninogenica* |
| *Streptococcus parasanguinis* clade 411 | *Streptococcus parasanguinis* |
| *Saccharibacteria* HMT352 | (excluded*) |

*No suitable representative identified within the AGORA metabolic models repertoire

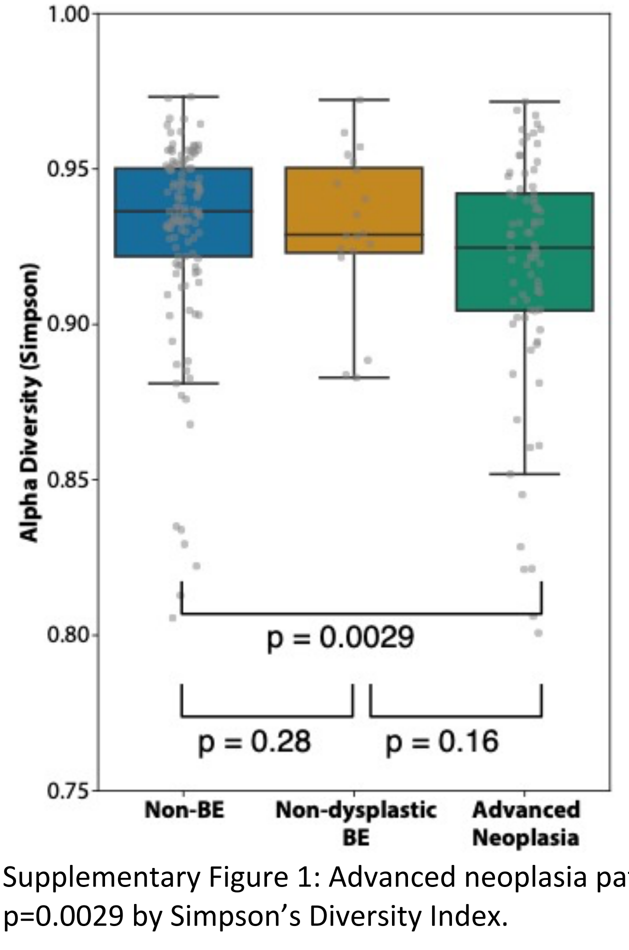


Supplementary Figure 1: Advanced neoplasia patients had significantly lower alpha diversity, p=0.0029 by Simpson's Diversity Index.

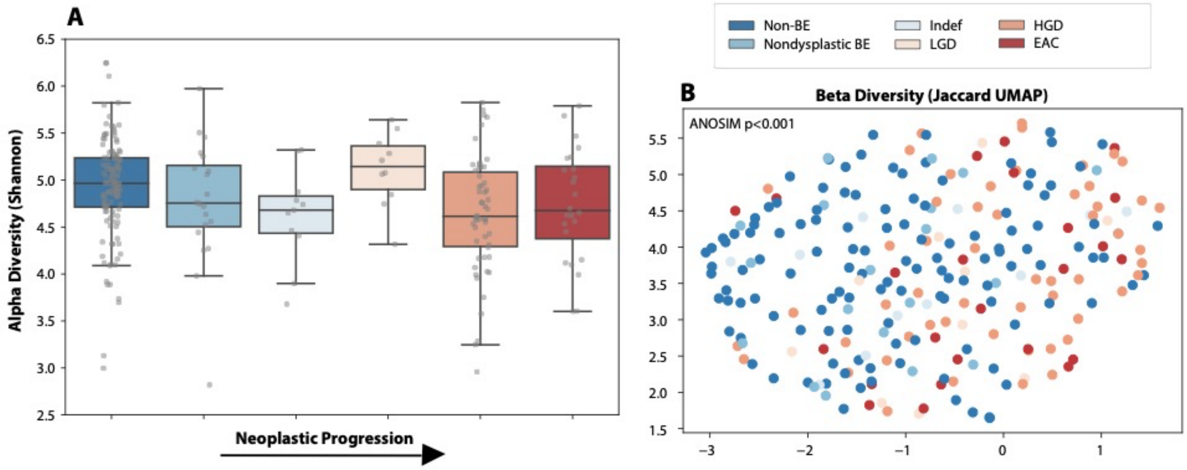


Supplementary Figure 2: (A) When separated into individual groups, there was a significant trend towards lower alpha diversity with stages of progression to EAC (p=0.006). (B) On Jaccard UMAP beta diversity analyses, there was clear visual clustering across stages of progression, and this was confirmed statistically (PERMANOVA p<0.001).

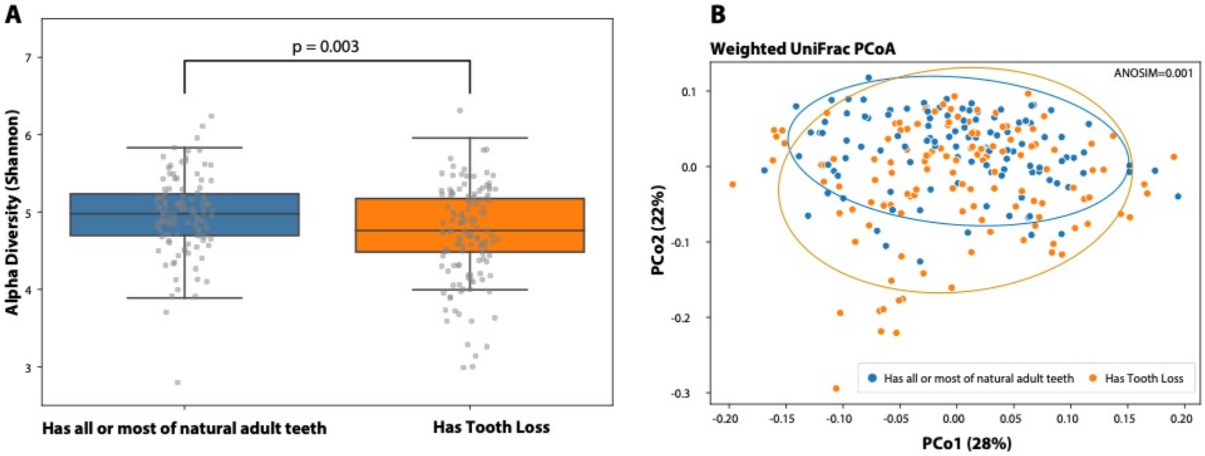


Supplementary Figure 3: Patients with tooth loss had significantly lower alpha diversity (p=0.003, Shannon; A) and had significantly different beta diversity (ANOSIM=0.001) on weighted UniFrac PCoA (B).

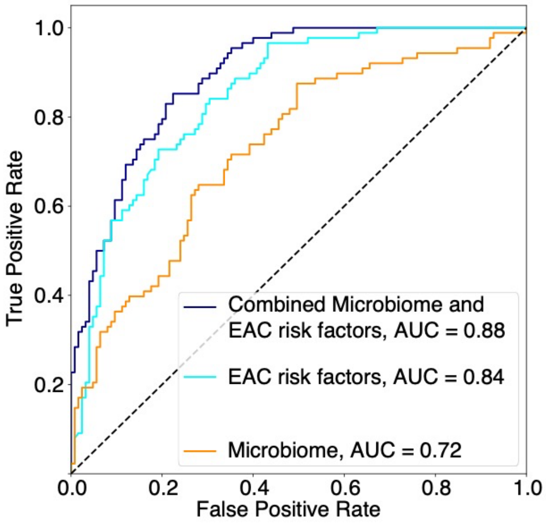


Supplementary Figure 4: A model to classify advanced neoplasia, trained on a combination of microbiome data and known EAC risk factors had a larger AUROC than a classifier model trained on known EAC risk factors alone, DeLong p=0.053.

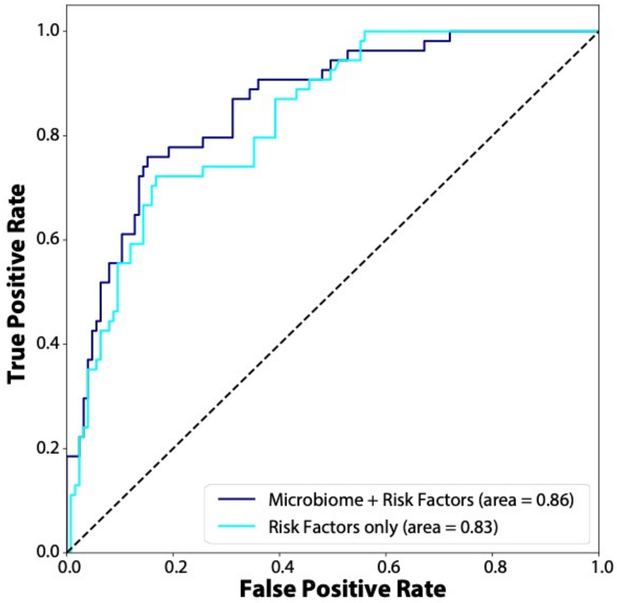


Supplementary Figure 5: A model to classify only high grade dysplasia vs non-BE showed and AUROC of 0.86 when both microbiome data and known clinical risk factors were included.
